# Cardiomyocyte GSK-3β deficiency induces cardiac progenitor cell proliferation in the ischemic heart through paracrine mechanisms

**DOI:** 10.1101/2021.08.28.458018

**Authors:** Ayesha M. Yusuf, Rizwan Qaisar, Abaher O. Al-Tamimi, Manju Nidagodu Jayakumar, James R. Woodgett, Walter J. Koch, Firdos Ahmad

## Abstract

Cardiomyopathy is an irreparable loss and novel strategies are needed to induce resident cardiac progenitor cell (CPC) proliferation *in situ* to enhance the possibility of cardiac regeneration. Here we sought to identify the potential roles of glycogen synthase kinase-3β (GSK-3β), a critical regulator of cell proliferation and differentiation, in CPC proliferation post-myocardial infarction (MI).

Cardiomyocyte-specific conditional GSK-3β knockout (cKO) and littermate control mice were employed and challenged with MI. Though cardiac left ventricular chamber dimension (LVID) and contractile functions were comparable at two-week post-MI, cKO mice displayed significantly preserved LV chamber and contractile function vs. control mice at four-weeks post-MI. Consistent with protective phenotypes, an increased percentage of c-kit-positive cells (KPCs) were observed in the cKO hearts at four and six-weeks post-MI which was accompanied by increased levels of cardiomyocyte proliferation. Further analysis revealed that the observed increased number of KPCs in the ischemic cKO hearts was mainly from a cardiac lineage as the majority of identified KPCs were negative for the hematopoietic lineage marker, CD45. Mechanistically, cardiomyocyte-GSK-3β profoundly suppresses the expression and secretion of growth factors, including basic-FGF angiopoietin-2, erythropoietin, stem cell factor (SCF), PDGF-BB, G-CSF, and VEGF, post-hypoxia.

In conclusion, our findings strongly suggest that loss of cardiomyocyte-GSK-3β promotes cardiomyocyte and resident CPC proliferation post-MI. The induction of cardiomyocyte and CPC proliferation in the ischemic cKO hearts is potentially regulated by autocrine and paracrine signaling governed by dysregulated growth factors post-MI. A strategy to inhibit cardiomyocyte GSK-3β could be helpful for the promotion of *in situ* cardiac regeneration post-ischemic injury.

## Introduction

Despite therapeutic advancement, adverse cardiac remodeling remains the leading cause of heart failure (HF) (Savarese and Lund, 2017). Acute myocardial infarction (AMI) often leads to maladaptive cardiac remodeling and is associated with cardiomyopathy and scar formation (Marian and Braunwald, 2017). Cardiac stress-induced cardiomyopathy and cardiac injury eventually promote interstitial fibrosis due to the inability of cardiac tissues to regenerate and extracellular matrix deposition (Laflamme and Murry, 2011; Lal et al., 2014; Porrello et al., 2011). The cardiac muscle is made up of a variety of cells which mainly comprise cardiomyocytes, fibroblasts, endothelial, and inflammatory cells. In addition to these, a limited number of cardiac progenitor cell (CPC) niches, with limited proliferation and differentiation capability, are also found in the adult heart (He et al., 2020). These cells in the heart facilitate normal cardiac contractile function through cell-to-cell direct communication and paracrine interaction (Yu and Wang, 2019).

MI and other cardiac stressor-induced cardiomyopathy and tissue damage represent an irreparable loss because of the minimal capacity of cardiomyocytes and CPCs to proliferate. Available therapeutic approaches are not efficient in limiting or reversing the maladaptive cardiac remodeling and chronic HF hence heart transplantation is the only and ultimate option. Unfortunately, heart transplantation is out of reach for the majority of HF patients. *In situ* cardiac regeneration has the potential to serve as a viable treatment option. Increasing evidence indicates that modulation of various cellular pathways, microRNAs, and exosomes are capable of inducing CPC proliferation in the injured heart (Iannolo et al., 2018; Witman et al., 2020; Zhang et al., 2014).

The α and β isoforms of glycogen synthase kinase-3 (GSK-3), a Ser/Thr kinase, regulate many physiological and pathophysiological processes through modulation of numerous cellular pathways (Ahmad and Woodgett, 2020; Lal et al., 2015). Several biological processes including glycogen synthesis, insulin resistance, mitochondrial function, cell proliferation, differentiation, and apoptosis, are regulated by GSK-3 isoforms (Ahmad et al., 2019; Beurel et al., 2015; Gupte et al., 2018). Most importantly, GSK-3 isoforms inhibit cardiomyocyte proliferation and cardiomyocyte-specific loss of GSK-3α or β promotes cardiomyocyte proliferation upon myocardial ischemia (Ahmad et al., 2014; Singh et al., 2019; Woulfe et al., 2010). Consistent with this finding, germline deletion of GSK-3β leads to hyper-proliferation of cardiomyoblast which results in obstructed ventricles and embryonic lethality of GSK-3β null mice (Kerkela et al., 2008). Based on these observations and other studies employing isoform-specific targeting of GSK-3 in embryonic stem (ES) cells, it is evident that GSK-3β is an essential regulator of stem cell proliferation and differentiation (Doble et al., 2007; Force and Woodgett, 2009). Interestingly, cardiomyocyte-specific loss of both GSK-3α and β isoforms promotes cardiomyocyte cell cycle reentry and mitotic catastrophe (Zhou et al., 2016a; Zhou et al., 2016b). These observations indicate that GSK-3β might have a similar effect on resident CPC proliferation in adult hearts as seen in cardiomyoblasts. As mentioned above, the limited number of resident CPCs in the adult heart restricts the capacity for cardiac regeneration. Therefore, targeting GSK-3β in adult cardiomyocytes would likely enhance resident CPC proliferation *in situ*.

Cardiomyocytes and other resident cardiac cells exert their regenerative effects primarily through paracrine mechanisms by releasing growth factors such as basic-fibroblast growth factor (bFGF), stem cell factor (SCF), vascular endothelial growth factor (VEGF), and hepatocyte growth factor (HGF) (Hodgkinson et al., 2016; Yoon, 2005). These paracrine factors facilitate cardiac regeneration through various mechanisms, including cytoprotection, induction of resident progenitor cell proliferation and differentiation, cardiomyocyte proliferation, and neovascularization. Emerging evidence suggests that paracrine factors have pleiotropic actions that help in cardiac repair. The release of paracrine factors may create a concentration gradient and therefore a tissue microenvironment that influences resident cell behavior leading to cardiac repair and regeneration (Hodgkinson et al., 2016).

GSK-3β regulates cardiomyocyte and other types of cell proliferation and differentiation (Force and Woodgett, 2009; Woulfe et al., 2010), however, the underlying mechanism is largely unknown. The growth factors including FGF, epidermal growth factor (EGF) and platelet-derived growth factor (PDGF) stimulate PI3K-Akt signaling and lead to Ser9 phosphorylation (inhibition) of GSK-3β (Cohen and Frame, 2001; Kaidanovich-Beilin and Woodgett, 2011; Takahashi-Yanaga, 2018). However, it is not clear whether dysregulated GSK-3β activity further modulates growth factor expression and secretion from cardiomyocytes under cardiac pathological conditions.

Here, we assess the potential roles of specific targeting of GSK-3β in the adult cardiomyocytes on resident CPC proliferation post-MI by employing a cardiomyocyte-specific conditional KO (cKO) mouse model. We report, for the first time, that loss of cardiomyocyte-GSK-3β promotes resident CPC proliferation post-MI which is accompanied by an increased degree of cardiomyocyte proliferation and discernible cardiac protection. Mechanistically, we identify that loss of cardiomyocyte-GSK-3β promotes cardiomyocyte and resident CPC proliferation, likely through crosstalk between GSK-3β-modulated cardiomyocytes and CPCs by modulating the levels of secreted growth factors, post-MI.

## Methods

### Generation of tamoxifen-inducible cardiomyocyte-specific GSK-3β knockout mouse

The cardiomyocyte-specific conditional GSK-3β knockout mouse model was generated as described previously (Woulfe et al., 2010). Briefly, mice carrying homozygous floxed alleles for exon 2 (flanked with lox-p sites) of GSK3B gene (Patel et al., 2008) were crossed with mice carrying α-myosin heavy chain (α-MHC) promoter-driven, tamoxifen-inducible *Mer-Cre-Mer* for two generations to generate *GSK3β*^*fl/fl Cre+/-*^ mice. The mouse line was maintained on the C57BL/6 background. The experimental animals were generated by crossing *GSK3β*^*fl/fl/ Cre+/-*^ *with GSK3β*^*fl/f*^ *mice*. Twelve-week-old male mice were treated with a tamoxifen chow diet for 15 days followed by two weeks on a regular chow diet to wash out the tamoxifen from their circulation. The customized tamoxifen chow diet containing 16% protein, 400mg/Kg tamoxifen citrate, 49.6g/Kg sucrose and 200mg/Kg red food color, was purchased from Teklad (Cat #TD. 130860). The tamoxifen-treated *GSK3β*^*fl/fl/ Cre+/-*^ *and GSK3β*^*fl/fl*^ *mice* were considered conditional knockout (cKO) and littermate controls, respectively. All animal procedures and treatments were performed as per *NIH guidelines and the protocol was* approved by the local Institutional Animal Care and Use Committee (IACUC) of Temple University.

### Myocardial infarction

Myocardial infarction or sham surgeries were performed as described previously (Gao, 2010). Briefly, cKO and littermate control mice were anesthetized using 2% isoflurane inhalation through an isoflurane vaporizing system (Viking Medical, Medford, NJ). A small skin incision (∼1.2cm) was made over the left chest and pectoralis major and minor muscles were dissected and retracted. The 4^th^ intercostal space was exposed and a small hole was made using a mosquito clamp to open the pleural membrane and pericardium. The heart was smoothly and gently “popped out” through the hole. The proximal left anterior descending (LAD) coronary artery was ligated with 6-0 silk suture. Once ligation was done, the heart was immediately placed back into the intra-thoracic space followed by manual air evacuation and closure of muscle and the skin. The mice were injected with *sustained*-release Buprenorphine SR-LAB (1.0 mg/kg) subcutaneous once after surgery or as needed.

### Echocardiography

Echocardiography was performed as described previously (Ahmad et al., 2020; Lal et al., 2012). Briefly, transthoracic two-dimensional motion mode-echocardiography was performed at 0, 2 and 4 weeks post-MI using a probe (VisualSonic, Vevo2100). Anesthesia was maintained throughout the echocardiography by supplying 1.5% isofluorane using a vaporizer. LV interior dimension at end-diastole (LVID;d) and end-systole (LVID;s) were measured, and ejection fraction (EF) and fractional shortening (FS) values were calculated using the Vevo2100 program.

### Histochemistry

The histochemistry was performed as described previously (Ahmad et al., 2020; Lal et al., 2012). Briefly, after 2, 4 and 8 weeks post-MI or sham surgeries, whole heart tissue was harvested from anesthetized mice and fixed in 4% paraformaldehyde followed by dehydration using increasing concentrations of ethanol, and then tissue was embedded in paraffin. Heart sections (5 μm) were stained using Masson’s trichrome kit (Sigma-Aldrich# HT15-1KT). The 0.8X and 20X objectives of a Nikon Eclipse 80i microscope were used to capture the whole heart and scar images and, images were analyzed for scar circumference, thickness and quantification of viable cardiomyocytes in scar using NIS Elements software.

### Immunofluorescence staining

Immunofluorescence staining of heart sections was performed as described previously (Ahmad et al., 2014). Briefly, heart sections were deparaffinized with xylene, rehydrated with successive incubations in decreasing ethanol concentrations, and finally incubated in distilled water. After antigen retrieval and 3X washing with phosphate-buffered saline (PBS), endogenous peroxidase activity was blocked by incubating the section with 0.15% hydrogen peroxide followed by blocking the sections with TNB (blocking) buffer (Perkin Elmer, Woodbridge, Ontario). Sections were incubated with primary antibodies [(c-Kit/CD117; R&D #AF1356; dilution 1:500), (CD45; Novus # NB100-77417; dilution 1:100), (Ki67; Vector Lab VP-RM04; dilution 1:100)] in TNB blocking buffer for overnight at 4°C followed by one-hour incubation with secondary antibodies: EnVisionTM^+^ System-HRP (DAKO K4002). TRITC-based Tyramide reagent pack from Perkin Elmer was used to amplify the fluorescence of Ki67 and c-Kit. Sections were then washed 3X with PBS and incubated with cardiomyocyte-specific primary antibody (Sarcomeric α-actinin; Sigma #A7811) for 60 min at 37°C followed by 3X wash with PBS. Slides were incubated with fluorescence-labeled secondary (Alexa fluor488) antibodies. After 3X washing with PBS, nuclei were stained by DAPI (1μg/ml) for 15 min. After a quick PBS wash, tissues were mounted with Vectashield (Vector Laboratories, Burlingame). Ki67 quantification was done by capturing five images from the border zone of LV using a Nikon Eclipse 80i microscope. A Carl Zeiss 510 confocal microscope was used to capture five images from the border zone of LV to analyze the c-Kit positive cells. All quantification and analyses were done using NIS Elements software.

### AC16 cell culture and plasmid transfection

AC16 human cardiomyocytes were purchased from EMD Millipore (SSC #109) and cultured in Dulbecco’s modified Eagle’s medium: Nutrient mixture F-12 (Sigma #D6434) supplemented with 2mM L-glutamine, 10% fetal bovine serum (FBS) and 1% Penicillin-Streptomycin antibiotic cocktail in a humidified 37°C incubator with 5% CO_2_. A total of 0.3 × 10^6^ AC16 cells were seeded onto each well of 6-well tissue culture plates. At ∼60% confluency, cells were serum-starved with 1X Opti-MEM (Gibco #00448) for one hour prior to transfection. Cells were transfected with a control Flag or HA-GSK-3β^WT human^ plasmid using Fugene 6 (Promega #E2693) in a 2:1 ratio and incubated for 3 hours in a CO_2_ incubator. After transfection, an equal volume of complete DMEM containing 2X FBS and 2X penicillin-streptomycin cocktail was added and incubated for 24 hours in a CO_2_ incubator.

### Heart and cell lysates preparation and western blotting

Cardiac tissue and cell lysate preparation, and Western blotting were done as described previously (Ahmad et al., 2020). Briefly, post-tamoxifen treatment cardiac tissues were excised from anesthetized animals and LV was homogenized using lysis buffer containing phosphatase and protease inhibitor cocktail. For AC16 cardiomyocytes, treated cells were washed with ice-chilled PBS twice and an appropriate amount of lysis buffer was added to each well of the culture plates. Cells were harvested using scrapers and lysates were collected in ice-chilled tubes. The homogenized LV and cell lysates were centrifuged at 15,000*g* for 15 min at 4°C. Supernatant was carefully transferred to fresh tubes and an equal amount of denatured protein was loaded onto SDS-PAGE. After trans-blotting, membranes were incubated with p-GSK-3α/β (Ser21/Ser9) (Cell Signaling #9331), GSK-3α/β (Cell Signaling #5676), GSK-3β (Cell Signaling #12456), or GAPDH (Fitzgerald # 10R-G109a) primary antibodies followed by incubation with appropriate HRP-labelled secondary antibodies. The blots were developed using enhanced chemiluminescence reagent (Bio-Rad #170-5060) and imaging was done under a gel documentation system (BioRad Chemidoc Touch Imaging System).

### Hypoxia stimulation and culture supernatant collection

A regulated Whitley H45 hypoxia chamber was used to simulate hypoxia in transfected AC16 cardiomyocytes. Prior to hypoxia, culture media was replaced with serum-free media, and cells were incubated for one hour in a CO_2_ incubator for acclimatization. Cells were then transferred to the hypoxia chamber containing 95% N_2_, 5% CO_2,_and 1% O_2_ concentrations which were maintained throughout the hypoxia period (24 hours). Once hypoxia was completed, culture plates were transferred to ice, and the culture supernatant was collected in ice-cold fresh tubes. The supernatant was centrifuged at 14,000 rpm for 5 minutes at 4°C to remove cell debris and the supernatant was transferred to a fresh tube and stored at −80°C until further use.

### Growth factor assay using flow cytometry-based LegendPlex kit

To assess levels of different growth factors including VEGF, basic-FGF (bFGF), stem cell factor (SCF), platelet-derived growth factor-AA (PDGF-AA), PDGF-BB, angiopoietin-2 (angpt2), epidermal growth factor (EGF), erythropoietin (EPO), hepatocyte growth factor (HGF), TGF-α, granulocyte colony-stimulating factor CSF (G-CSF), macrophage CSF (M-CSF), and granulocyte-macrophage CSF (GM-CSF) in the culture supernatant and cell lysates, a multiplex bead-based LegendPlex assay (BioLegend #740180) was performed following the manufacturer’s instruction. Briefly, culture supernatant was mixed with an equal volume of assay buffer or equal amount of protein lysate (10µg) was diluted in assay buffer and loaded onto an assay plate and beads were added to each well. The sealed plate was incubated on a plate-shaker at 300 rpm for 2 hours at room temperature. The assay plate was centrifuged, and the solution aspirated. Assay beads were washed with wash buffer twice and detection antibodies were added to each well. The sealed plate was incubated on a plate-shaker at 300 rpm for 1 hour at room temperature. SA-PE solution was added to each well and incubated on a shaker for another 30 minutes. The supernatant was carefully aspirated from each well and beads were re-suspended in wash buffer. Data were acquired using a BD FACS Aria III with FACS Diva software, and data analysis was done using LegendPlex software (Biolegend, USA). All samples were assessed in duplicate and the average was calculated.

### Statistics

Two-way ANOVA followed by Tukey’s post hoc test for multiple comparisons or unpaired *t*-test for single comparison (Graph Pad Prism Software Inc., San Diego, CA) was performed to evaluate the significant differences between data groups. Data are expressed as mean ± SEM. For all tests, a *p*-value <.05 was considered for statistical significance.

## Results

### Loss of Cardiomyocyte-GSK-3β limits LV chamber dilatation and contractile dysfunction post-MI

Given the critical role of GSK-3β in cell proliferation and differentiation (Ji et al., 2015; Korur et al., 2009), we hypothesized that GSK-3β regulate cardiac progenitor cell proliferation in the ischemic heart. In this series, 12-14 week old cKO and littermate control mice were treated with tamoxifen and GSK-3β protein expression was assessed in the heart by immunoblotting. GSK-3β protein expression levels were reduced by ∼80% in the cKO vs. control LV lysates post-tamoxifen treatment (**Fig. 1A-B**). The residual GSK-3β protein (∼20%) in the cKO was most likely derived from other types of cardiac cells including fibroblast and endothelial cells.

**Figure 1.**
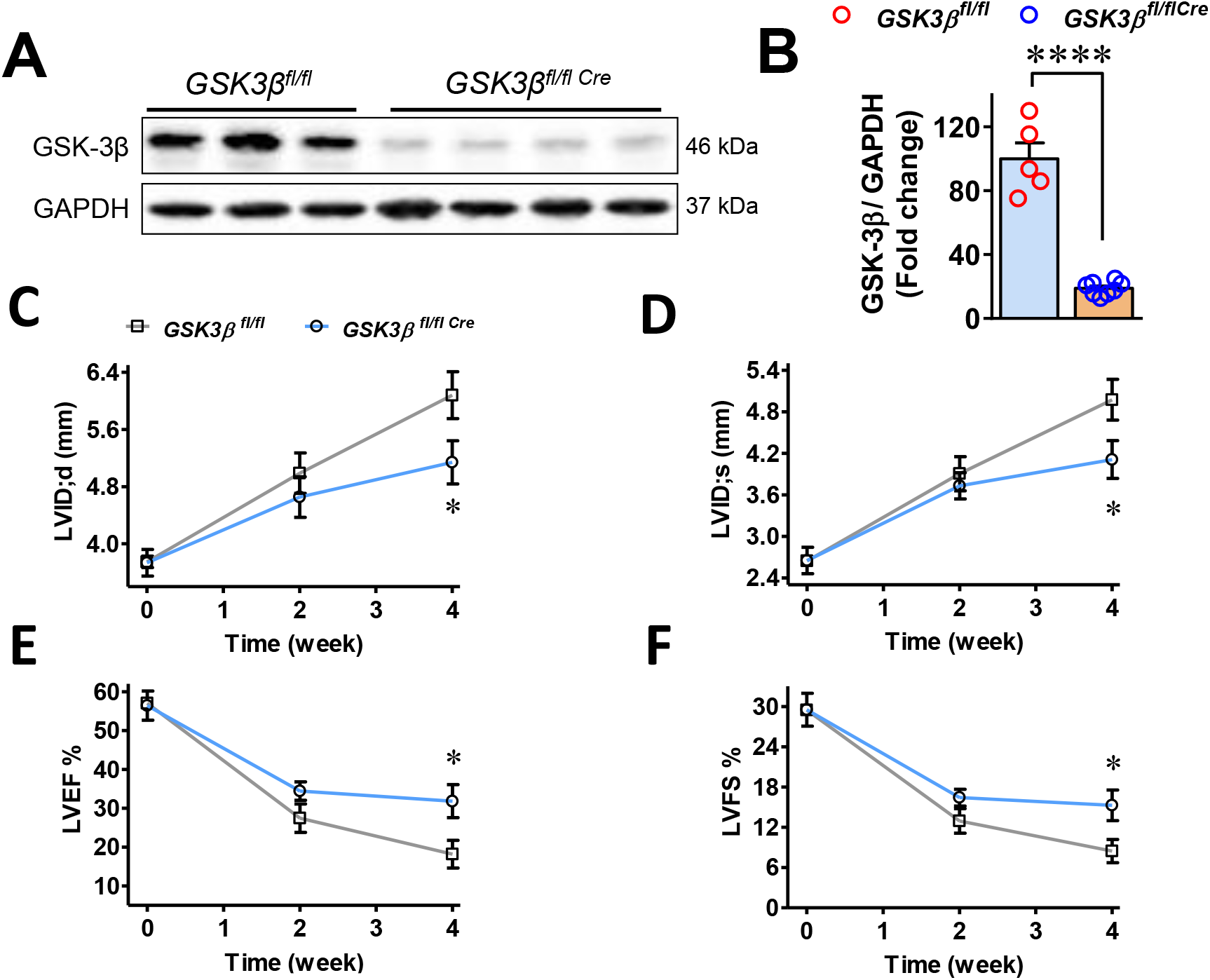
Conditional genetic ablation of *GSK-3β* attenuates post-MI adverse cardiac remodeling and dysfunction: Post-tamoxifen treatment, *GSK3β*^*fl/flCre*^ and littermate controls *GSK3β*^*fl/fl*^ were assessed for GSK-3β protein expression. (**A**) representative blot image and (**B**) quantification show ∼80% reduction in GSK-3β expression in *GSK3β*^*fl/flCre*^ (cKO) in comparison to *GSK3β*^*fl/fl*^ (control) hearts post-tamoxifen treatment. An unpaired *t*-test was performed to compare the group. n=5–8; ^****^ *P*< 0.0001. The left ventricular interior chamber dimension (LVID) both at (**C**) end-diastolic (LVID;d) and (**D**) end-systolic (LVID;s) were comparable at 0 and 2 weeks post-MI. However, comparatively smaller LV chambers in the cKO animals both at end-diastolic and systolic at four week post-MI show preserved LV chamber in the cKO post-MI. (**E**) Traces show a comparable LV ejection fraction (LVEF) and (**F**) fractional shortening (LVFS) at 0 and 2 weeks post-MI and significantly preserved contractile functions in the cKO vs. control animals at four weeks post-MI. An unpaired *t*-test was performed to compare the group. n=6–13; ^*^ *P*< 0.05.

Next, post-tamoxifen treatment, a batch of cKO and littermate control mice was recruited and MI surgery was performed through LAD ligation. Cardiac function was assessed through serial two-dimensional motion-mode echocardiography at 0, 2 and 4 week time points. At baseline, the left ventricular (LV) chamber and cardiac contractile functions were comparable between cKO and littermate controls. The LV chamber dilated and contractile functions, both at systole and diastole, deteriorated equally both in the cKO and control mice up to 2 weeks post-MI. At four week post-MI, LV chamber both at systole and diastole were significantly better preserved in the cKO compared to control mice (**Fig. 1C-D**). The preserved LV chamber dimensions in the cKO were reflected in contractile function where LV ejection fraction (LVEF) and fractional shortening (LVFS) were significantly better in the cKO vs. control mice (**Fig. 1E-F**). These data strongly suggest that targeting specifically GSK-3β in adult cardiomyocytes protects against pathological cardiac remodeling and preserves cardiac contractile functions post-MI.

### Cardiomyocyte-GSK-3β deficiency promotes cardiomyocyte and cardiac progenitor cell proliferation post-MI

This study was designed to assess the role of GSK-3β in cardiac stem cell proliferation. Therefore, cardiac tissues were harvested from anesthetized cKO and control mice after 2, 4 and 8 weeks post-MI. Cardiac tissues from all three groups were processed for histochemistry and immunofluorescence staining. First, cardiomyocyte proliferation was assessed in 4-week post-MI cKO hearts through Ki67 (a mitosis marker), along with α-actinin (a cardiomyocyte-specific marker) staining. Only cardiomyocytes, positive for Ki67 co-localizing with 4,6-diamidino-2-phenylindole (DAPI), were counted and considered to be proliferating cardiomyocytes. A significantly higher number of Ki67 positive cardiomyocytes was observed in the cKO vs. control hearts post-MI (**Fig. 2A-B**).

**Figure 2.**
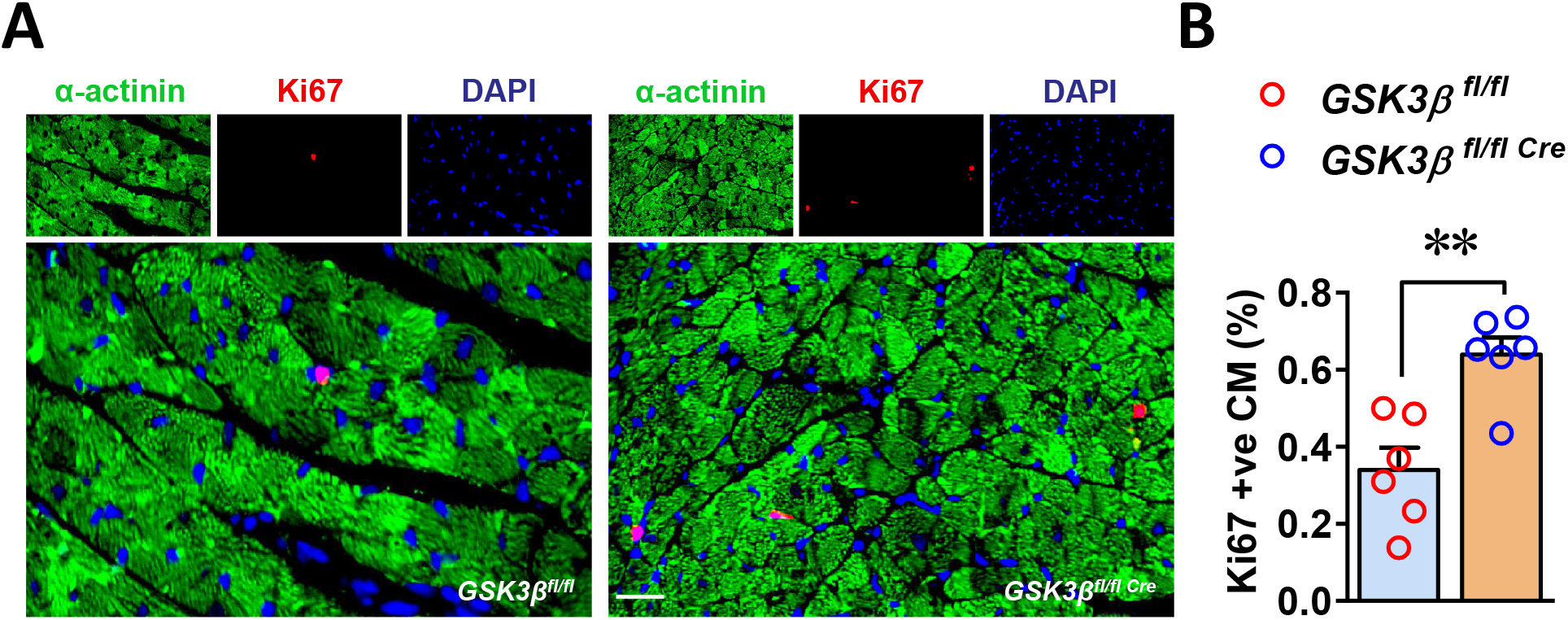
Loss of cardiomyocyte-GSK-3β promotes cardiomyocyte proliferation post-MI. **(A)** Representative images of Ki67 positive cardiomyocytes from GSK-3β cKO and control heart sections from four week post-MI cardiac tissue. (**B**) Bar diagram from Ki67-positive cardiomyocytes quantification shows a significantly increased number of Ki67-positive cardiomyocytes in the cKO vs. control hearts post-MI. An unpaired *t*-test was performed to compare the group. ^**^ *P*<0.005.

Next, we investigated whether GSK-3β deficiency in cardiomyocytes also impact resident CPC proliferation during ischemic conditions. To address the same, the heart sections from 2, 4 and 8 week post-MI animal groups were immunostained for c-Kit, a CPC-specific marker, along with α-actinin. The quantification of c-Kit-positive cells (KPC) in the cKO and control hearts revealed a comparable number of KPCs in 2-week post-MI cKO and control hearts; however, surprisingly, the number of KPCs was significantly higher in the cKO vs. control hearts at 4-week post-MI. The increased number of KPCs was sustained in the cKO hearts at 8-week post-MI (**Fig. 3A-B**). These findings strongly suggest that GSK-3β deficiency in cardiomyocytes promotes both cardiomyocyte and KPC proliferation in the ischemic heart.

**Figure 3.**
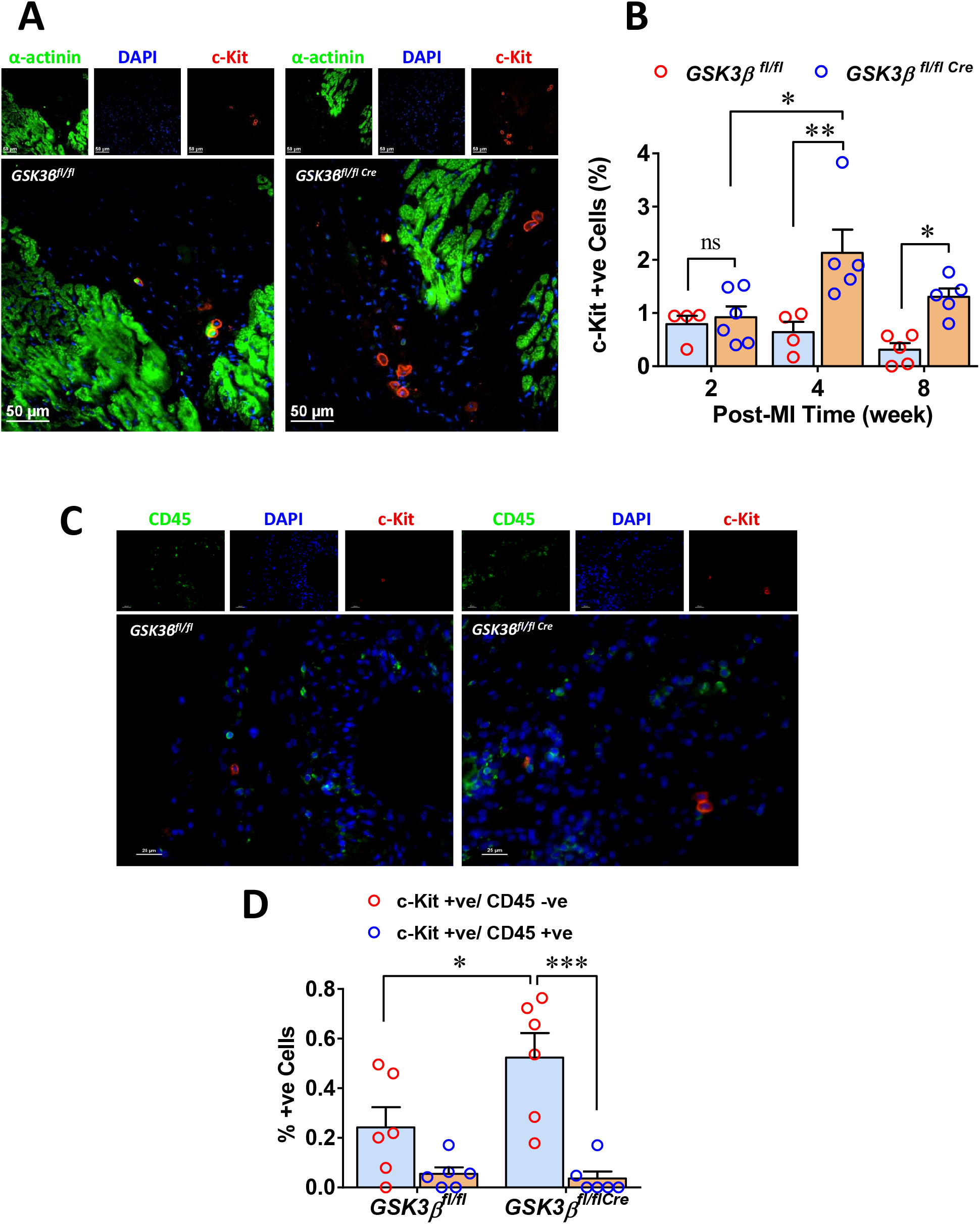
Cardiomyocyte-GSK-3β deficiency promotes cardiac progenitor cell (c-Kit cell) proliferation post-MI. (**A**) Representative confocal images show c-Kit–positive cells in ischemic cKO and control hearts. (**B**) c-Kit-positive cell quantification shows comparable numbers in the cKO vs. control hearts at 2 weeks and a significantly increased percentage of c-Kit-positive cells at 4 and 8 weeks post-MI. (**C**) Representative immunofluorescence staining images show CD45 and c-Kit positive cells. (**D**) Quantification revealed that the majority of c-Kit-positive cells identified in the ischemic hearts were CD45 negative. A two-way ANOVA followed by Tukey’s multiple comparisons was performed to compare the group. n=4-6; ^*^ *P*<0.05,^**^*P*<0.005^***^ *P*<0.001.

### The increased number of c-Kit positive cells in the ischemic cKO heart is contributed specifically due to resident CPC proliferation

To assess if the observed elevated number of KPCs in the ischemic cKO hearts was due to resident CPC proliferation or to increased infiltration of hematopoietic lineage cells in the cKO hearts post-MI, 4-week post-MI heart sections (as the changes in c-Kit positive cells were maximum at this time point) were immuno-stained for c-Kit along with CD45, a hematopoietic lineage marker. Cells positive for c-Kit, and dual positive cells for both c-Kit along with CD45 were quantified. Indeed, the percentage of dual positive cells was significantly lower than cells positive only for c-Kit, both in cKO and control hearts, post-MI. Consistent with previous experiments, a significantly higher number of KPCs was observed in the cKO vs. control hearts post-MI (**Fig. 3C-D**). These results attest that the observed increased number of KPCs in post-MI cKO hearts was derived from resident CPC proliferation.

### GSK-3β promotes scar expansion and thinning post-MI

Since a better preserved LV chamber dimension and an enhanced level of cardiomyocyte and resident CPC proliferation was observed in the cKO heart post-MI, we next sought to assess the LV chamber and characteristics of scar in the cKO hearts. Masson’s trichrome staining followed by imaging of whole heart sections revealed a well-expanded scar in both cKO and control hearts (**Fig. 4A**). Although there was a trend towards attenuated scar formation in the cKO vs. control hearts, it was not significant (**Fig. 4B**). Interestingly, scar thickness was much greater in the cKO hearts (**Fig. 4C**). Consistently, a significantly higher number of viable cardiomyocytes was observed in the scar area of cKO hearts (**Fig. 4D-E**). These observations suggest that, consistent with increased cardiomyocyte and CPC proliferation post-MI, deletion of cardiomyocyte-GSK-3β limits LV chamber dilatation and attenuates scar thinning post-MI.

**Figure 4.**
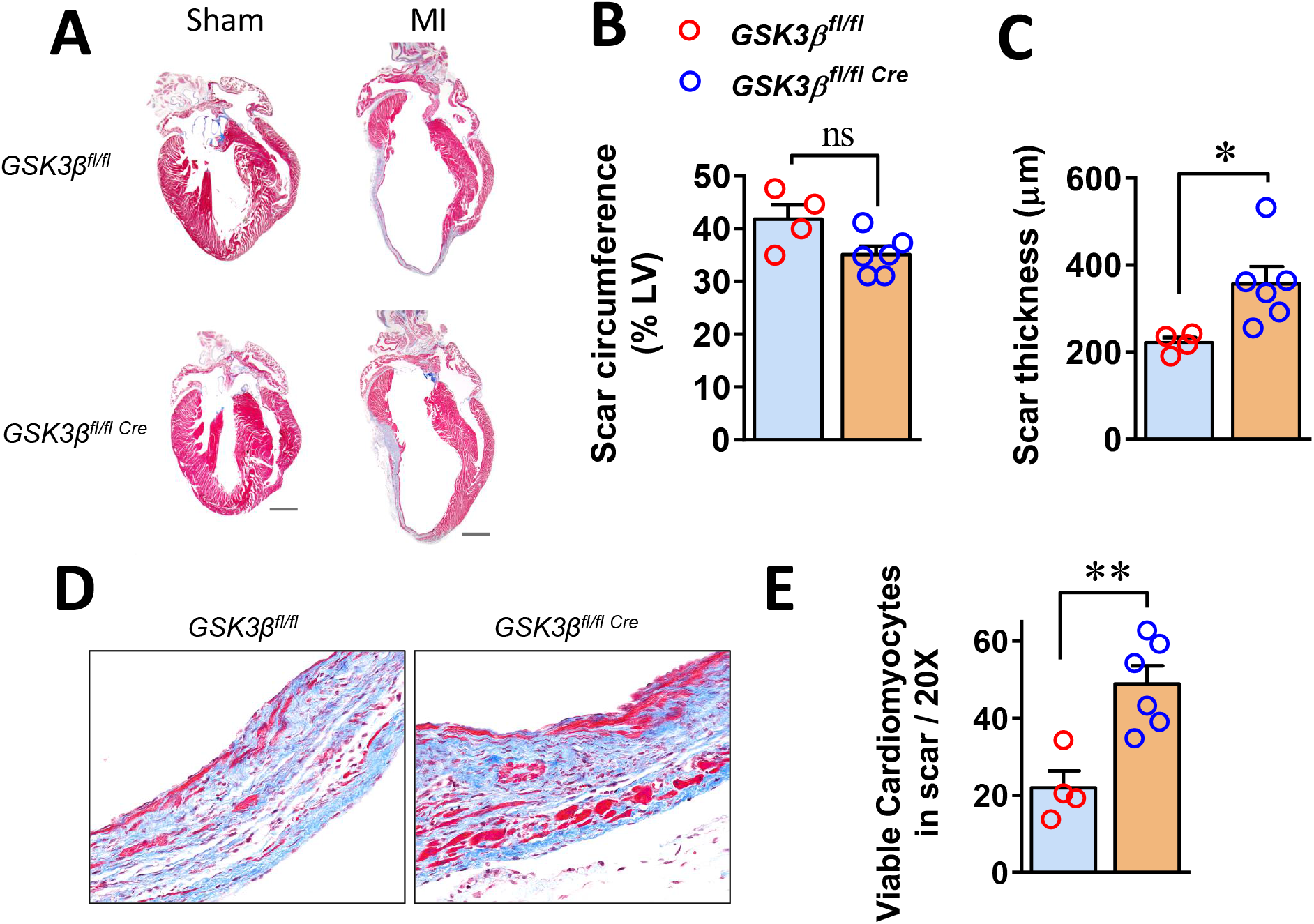
Loss of cardiomyocyte-GSK-3β attenuates scar formation post-MI: (**A**) Representative images show Masson’s trichrome-stained heart sections from sham or four-week MI operated cKO and littermate control hearts. (**B**) Bar diagram shows scar thickness in the cKO and control hearts. (**C**) The bar diagram shows a significantly smaller scar size in the cKO vs. control hearts. (**D**) Representative images taken under 20X objective show viable cardiomyocytes (red) in the scar area of control and cKO hearts. (**E**) Quantification shows a significantly increased number of viable cardiomyocytes in the scar area of cKO vs. control hearts. An unpaired *t*-test was performed to compare the group. ns, non-significant; ^*^ *P*<0.05.

### Cardiomyocyte-GSK-3β dysregulates hypoxia-induced growth factor expression and secretion from cardiomyocytes

Though the elevated levels of cardiomyocytes and resident CPC proliferation were observed in the cKO hearts post-MI, it is not clear how, specifically, suppression of GSK-3β function in the cardiomyocytes induces the resident CPC to proliferate post-MI. To address this question, we assessed the possible paracrine signaling mechanisms governed by numerous growth factors secreted by cardiomyocytes, post-MI. In this series, first we assess the level of GSK-3β activity in the cardiomyocytes post-hypoxia through assessing the phosphorylation levels of Ser9. The level of GSK-3β phosphorylation (inhibition) was significantly higher in the cardiomyocytes post-hypoxia (**Fig. 5A-B**). Next, we employed a GSK-3β gain-of-function model in AC16 cardiomyocytes. First, the level of GSK-3β protein expression was assessed in the cardiomyocytes engineered to overexpress the protein kinase. GSK-3β protein expression was ∼4 fold higher in the HA-GSK-3β plasmid transfected compared to flag transfected cells. The level of GSK-3α protein was comparable in both groups (**Fig. 5C-D**).

**Figure 5.**
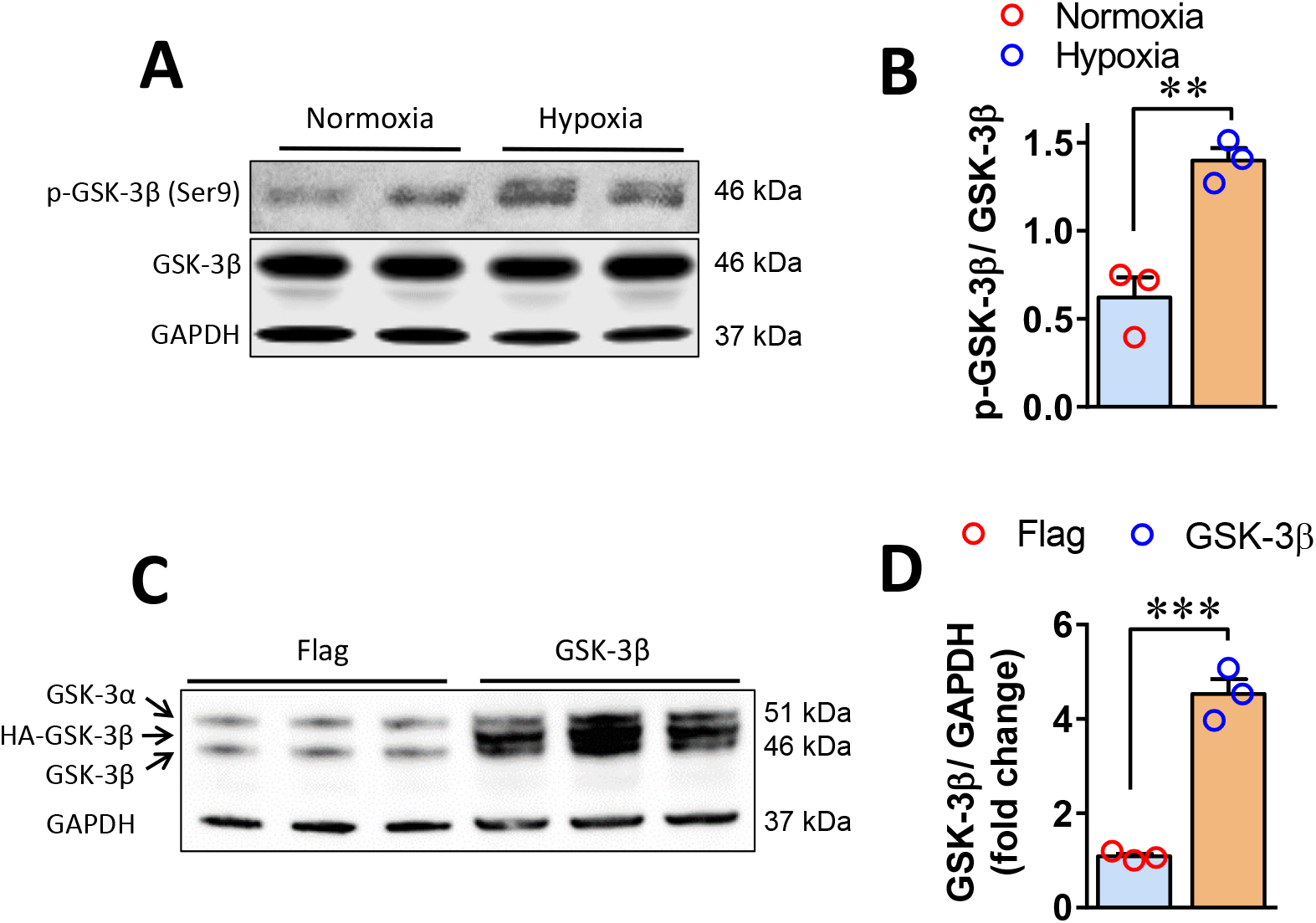
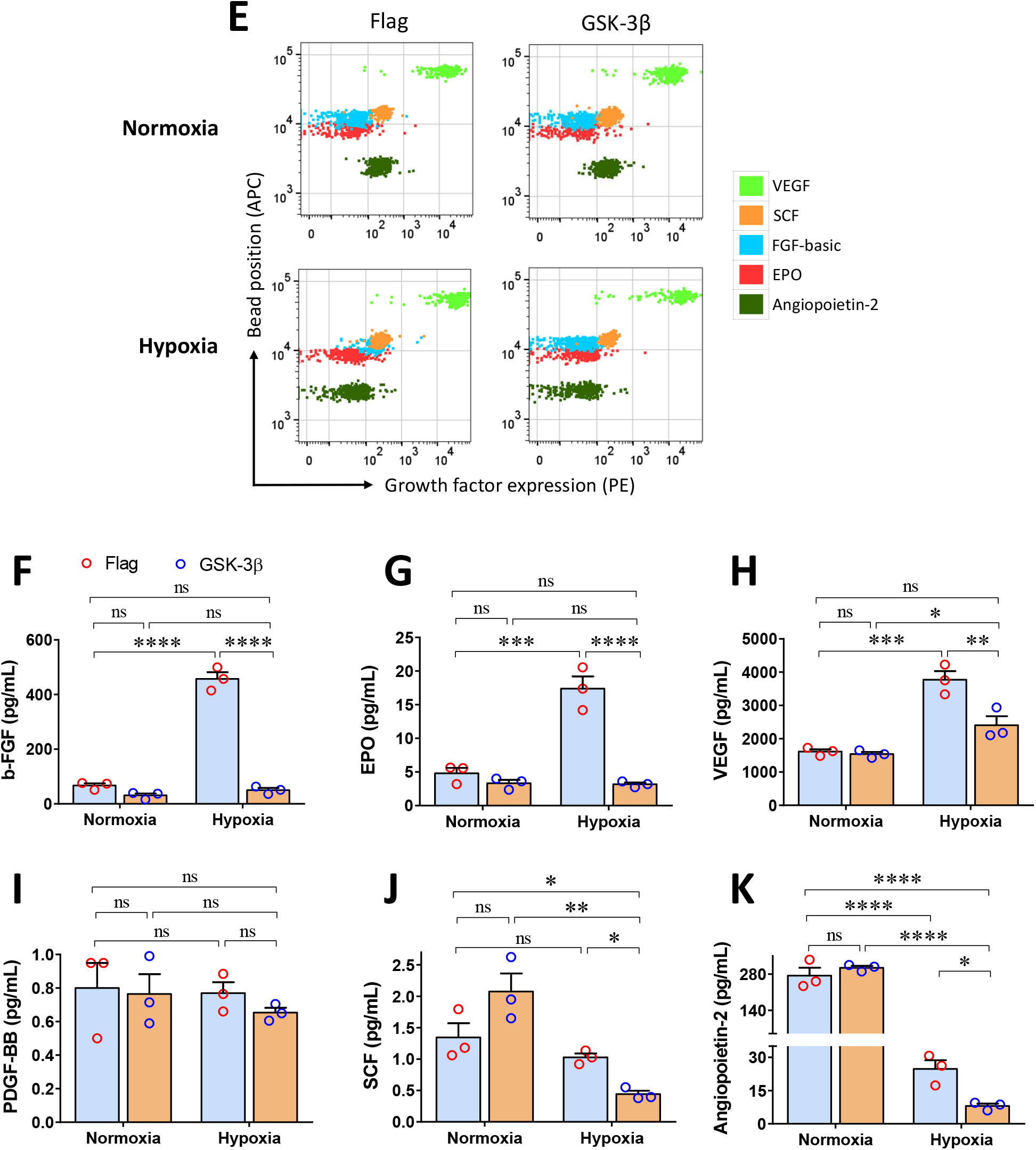
GSK-3β suppresses growth factor expression in cardiomyocytes post-hypoxia: (**A**) Representative blot images and (**B**) quantification show a significantly increased phosphorylation (inhibition) of GSK-3β in AC16 cardiomyocytes post-24 hours hypoxia. (**C**) The blot image and (**D**) quantification show ∼4 fold increased expression of GSK-3β in AC16 cardiomyocytes transfected with HA-GSK-3β (human) plasmid. (**E**) Representative flow cytometry scatter dot plots show the levels of growth factors in the supernatant of Flag vs. HA-GSK-3β transfected cardiomyocytes under normoxia and hypoxia conditions. Bar diagrams show a significantly lower level of (**F**) basic-fibroblast growth factor (b-FGF), (**G**) erythropoietin (EPO), (**H**) vascular endothelial growth factor (VEGF), (**I**) platelet-derive growth factor-BB (PDGF-BB), (**J**) stem cell factor (SCF), and (**K**) the level of angiopoietin-2 in the culture supernatant of GSK-3β overexpressing cardiomyocyte post-hypoxia. A two-way ANOVA followed by Tukey’s multiple comparisons was performed to compare the group. ns, non-significant; ^*^ *P*<0.05, ^**^ *P*<0.005, ^***^*P*<0.001, ^****^ *P*<0.0001.

After characterization of GSK-3β protein expression, AC16 cells were challenged with chronic hypoxia and culture supernatant was assessed for the level of several important human growth factors to identify potential autocrine and paracrine roles of GSK-3β in the cardiomyocyte and CPC proliferation. The levels of growth factors bFGF, EPO, VEGF, PDGF-BB, SCF, and angpt2 were comparable between the control and GSK-3β overexpressing cells under normoxia conditions. Interestingly, induction of particularly bFGF, EPO, and VEGF was observed in the control group post-hypoxia, which was significantly attenuated in GSK-3β overexpressing group (**Fig. 5E-H**). The level of PDGF-BB was unchanged in control groups subjected to normoxia or hypoxia, however; a slightly lower level was detected in the GSK-3β overexpressing vs. control cells challenged with hypoxia (**Fig. 5I**). Similarly, the level of SCF was found comparable in the normoxia groups and a significantly lower level was identified in the GSK-3β overexpressing in comparison to the control cells, post-hypoxia (**Fig. 5J**). Moreover, irrespective of levels of GSK-3β gene expression in the cardiomyocytes, the level of angpt2 was dramatically decreased in the hypoxia vs. normoxia group.

However, the level was even lower in GSK-3β overexpressing versus control cells that had been exposed to hypoxia (**Fig. 5K**). The levels of PDGF-AA and TGF-α were unchanged in both cell groups subjected to hypoxia and, EGF and HGF were undetectable in the culture supernatant (**Supplemental Fig. 1**). These findings suggest that GSK-3β is compensatorily inhibited in ischemia conditions. Moreover, data indicate that GSK-3β is a critical regulator of major growth factors in cardiomyocytes and that GSK-3β overexpression inhibits hypoxia-induced growth factor expression and secretion from cardiomyocytes.

Next, we assess the levels of the aforementioned growth factors in the cell lysates to assess the level of expression and for potential secretion defects. The comparable levels of b-FGF, EPO, VEGF, PDGF-BB, SCF, angpt2, TGF-α, G-CSF and M-CSF were identified in the GSK-3β overexpressing and control cardiomyocytes post-hypoxia (**Supplemental Fig. 2-3**). The level of PDGF-AA was upregulated, whereas decreased levels of G-CSF, M-CSF and GM-CSF were found in the GSK-3β overexpressing cardiomyocytes post-hypoxia (**Supplemental Fig. 3**). These findings support the idea that GSK-3β promotes the expression of PDGF-AA in cardiomyocytes post-hypoxia. However, the levels of this growth factor were unchanged in the culture supernatant because the secretion was not induced.

### GSK-3β overexpression inhibits CSFs in cardiomyocytes post-hypoxia

Colony-stimulating factors (CSFs) are another class of glycoproteins that bind to receptors present on stem cells and induce intracellular signaling pathways which promote cell proliferation and differentiation. Analysis of culture supernatant revealed a differential secretion of key CSFs from GSK-3β overexpressing cardiomyocytes. The level of G-CSF was comparable between control and GSK-3β overexpressing groups subjected to normoxia, however; the levels were significantly blunted in culture supernatant from GSK-3β overexpressing cells post-hypoxia (**Fig. 6A-B**). In contrast, M-CSF was comparable in the hypoxia groups and an elevated level was identified in the GSK-3β overexpressing cells under normoxia (**Fig. 6C**). A minimal effect of hypoxia was seen on levels of GM-CSF as a comparable amount was detected in normoxia vs. hypoxia groups irrespective of the level of GSK-3β. However, a significantly decreased level of GM-CSF was observed in GSK-3β overexpressing compared to the control group under both normoxia and hypoxia conditions (**Fig. 6D**). These findings suggest that GSK-3β regulates the expression of M-CSF in cardiomyocytes, particularly under hypoxia conditions, and regulates GM-CSF under both normoxia and hypoxia.

**Figure 6.**
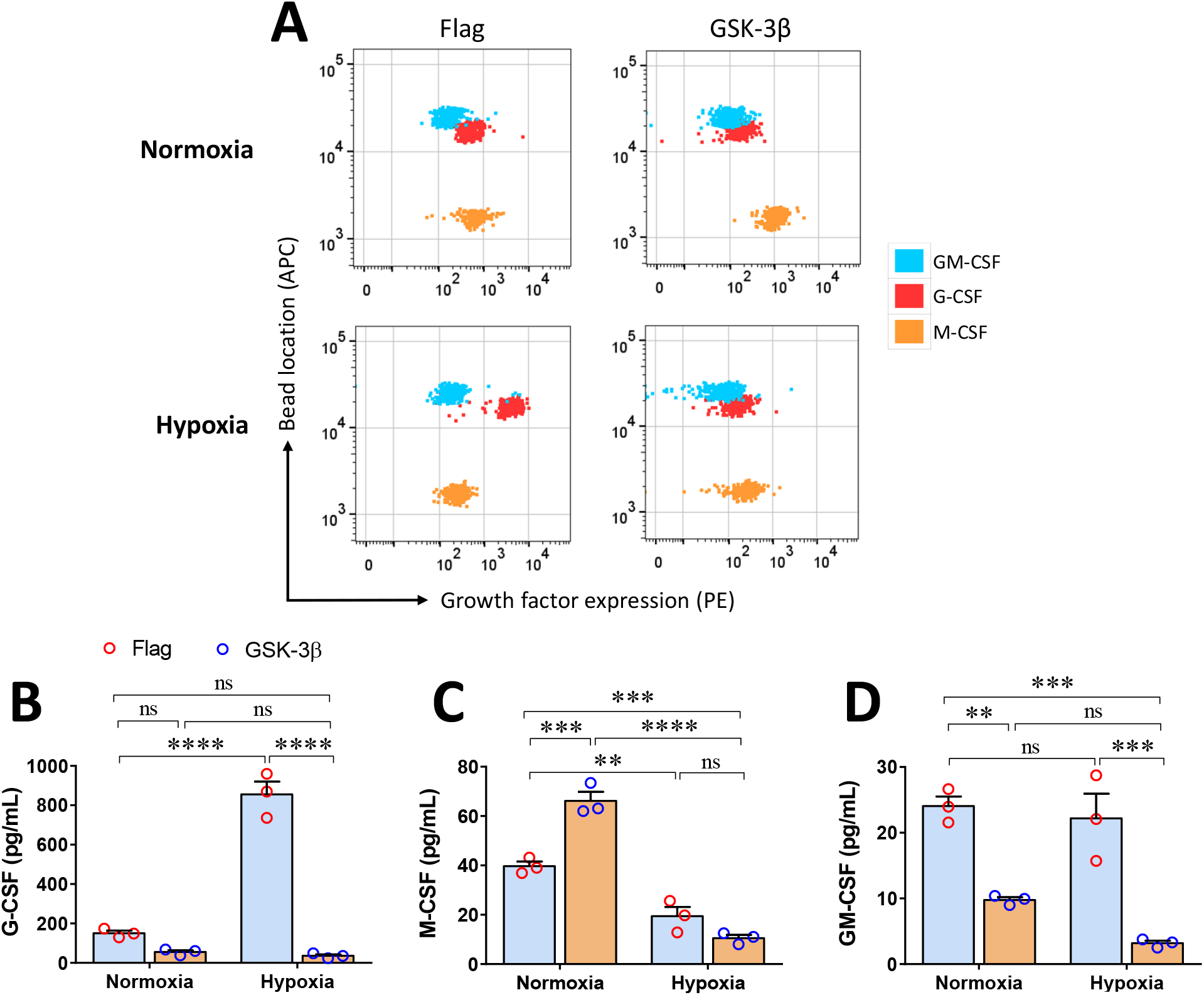
GSK-3β regulates colony-stimulating factors in the cardiomyocytes post-hypoxia: (**A**) Representative flow cytometry dot plots show the levels of different colony-stimulating factors in the the culture supernatant of Flag vs. HA-GSK-3β transfected cardiomyocytes under normoxia and hypoxia conditions. Bar diagrams show (**B**) lower level of granulocyte colony-stimulating factors (G-CSF), (**C**) comparable macrophage colony-stimulating factors (M-CSF), and (**D**) lower level of granulocyte-macrophage colony-stimulating factors (GM-CSF) in the supernatant of GSK-3β overexpressing cardiomyocytes vs. control cells post-hypoxia. A two-way ANOVA followed by Tukey’s multiple comparisons was performed to compare the group. ns, non-significant; ^**^ *P*<0.005, ^***^ *P*<0.001, ^****^ *P*<0.0001.

## Discussion

Cardiac endogenous regenerative capacity is limited and largely unable to replace cardiac tissue post-injury due to the presence of a limited number of resident CPCs niches and, the non-dividing nature of cardiomyocytes (Ahuja et al., 2007; Segers and Lee, 2008). Strategies like *ex vivo* expansion of CPCs followed by injection, infusion, or transplantation have been attempted to regenerate injured cardiac tissue. In this study, employing the cardiomyocyte-specific cKO mouse model we show, for the first time, that cardiomyocyte-GSK-3β acts to suppress resident CPC proliferation and conditional deletion leads to CPC and cardiomyocyte proliferation, post-MI. These observations were accompanied by a protective phenotype including protection of contractile function and preservation of LV chamber dimension in the cKO mice. Histological assessment revealed a comparatively thicker scar with a significantly increased number of viable cardiomyocytes in the cKO compared to controls, post-MI. Moreover, we identified for the first time, that GSK-3β regulates the expression and secretion of specific growth factors, under stress conditions, known to regulate cardiac regeneration (**Fig. 7**). These findings suggest that CPC and adult cardiomyocyte proliferation in cKO hearts occurs due to paracrine and autocrine mechanisms.

**Figure 7.**
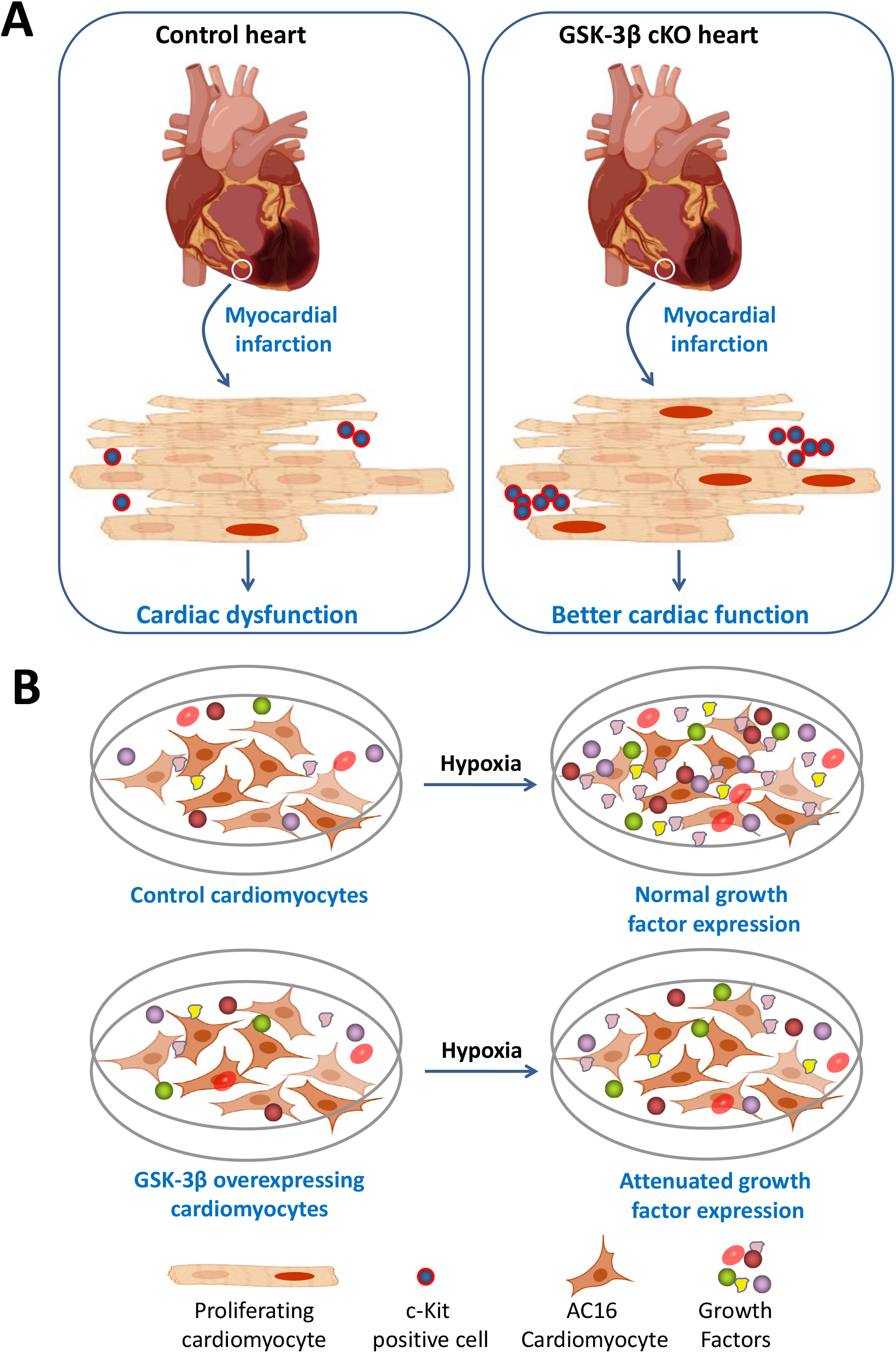
Cardiomycocyte-GSK-3β regulates CPCs and cardiomyocyte proliferation post-MI: **(A)** Schematic diagram shows that GSK-3β inhibits proliferation of cardiomyocyte and c-kit –positive cell (cardiac progenitor cell), and cardiomyocyte-specific deletion of GSK-3β (cKO) promotes proliferation of these cardiac cells post-myocardial infarction (MI). (**B**) Representative diagram shows that the gain-of- GSK-3β function in the cardiomyocytes highly suppresses the expression and secretion of numerous growth factors including VEGF, basic-FGF (bFGF), erythropoietin (EPO), stem cell factor (SCF), platelet-derived growth factors-BB (PDGF-BB), angiopoietin-2, granulocyte colony-stimulating factor CSF (G-CSF), and granulocyte-macrophage CSF (GM-CSF) under hypoxia, those are essentially required for cell proliferation, differentiation and survival. The findings strongly indicate that cardiomyocyte-GSK-3β regulates cardiomyocyte and cardiac progenitor cell proliferation post-MI potentially through autocrine and paracrine signaling mechanisms.

The role of GSK-3β in embryonic cardiomyoblast proliferation and differentiation was previously established and was linked to increased expression of cell cycle regulators. One study reported that germline deletion of GSK-3β causes neonatal death due to the thickening of LV and RV walls, which were filled with proliferating immature cardiac cells (Kerkela et al., 2008). Moreover, conditional deletion of GSK-3β enhances the proliferative capacity of adult cardiomyocytes, but only after MI (Woulfe et al., 2010). Our study confirms and extends these findings and shows that conditional loss of GSK-3β promotes not only adult cardiomyocytes but also CPC niche activation and proliferation, post-MI. A noticeably increased number of viable cardiomyocytes in the scar of cKO animals likely contributed due to cardiomyocyte proliferation, which can only be confirmed through lineage tracing. The viable cardiomyocytes in the scar may provide a better cardiac force-generating capacity resulting in better cardiac contractile function in the cKO mice post-MI.

Our findings indicate an interplay of autocrine and paracrine mechanisms for cardiac regeneration in the cKO mice, post-MI. The autocrine effects involve secreted factors that potentially induce the proliferation of cardiomyocytes. The paracrine effects involve the activation of neighboring cells, mainly the CPC niches, through factors secreted by cardiomyocytes. Our *in vitro* studies with the GSK-3β gain-of-function model have unraveled the dysregulation of a number of candidate growth factors possibly involved in cardiac restoration through autocrine and/or paracrine effects. It is evident that ligand binding with growth factor (GF) receptors present on cardiomyocytes and CPCs leads to their activation and proliferation (Urbanek et al., 2005a; Urbanek et al., 2005b). Previous studies have shown that SCF, a c-Kit ligand, promotes chemotactic properties of progenitor cells though the level of SCF was downregulated post-MI (Chute et al., 2006; Woldbaek et al., 2002). Consistently, we identified decreased SCF levels in the hypoxic compared to normoxic cardiomyocytes and an increased number of c-Kit positive cells in the cKO heart, post-MI. Moreover, a comparatively lower level of SCF, particularly in GSK-3β overexpressing cells undergoing hypoxia, indicates that GSK-3β further suppresses SCF expression, post-MI. SCF has limited efficacy alone, but when mixed with other growth factors such as GM-CSF and EPO, it displays a synergistic effect, particularly on hematopoietic stem cells (Bernstein et al., 1991; Williams et al., 1992). We observed that GSK-3β attenuates the levels of both GM-CSF and EPO, along with SCF, in cardiomyocytes post-hypoxia and, observed increased number of CPCs in the cKO post-MI in this study is a possible synergistic paracrine effect of different growth factors.

Hematopoietic stem cells also play important roles in cardiac restoration following injury (Der Sarkissian et al., 2017), however, we did not find a significant infiltration of hematopoietic stem cells in the cKO heart, post-MI. Thus, we conclude that loss of cardiomyocyte-GSK-3β elevates the levels of growth factors post-MI and creates a local niche, which activates resident CPCs, independent of hematopoietic stem cells. There is also a possibility that targeting cardiomyocyte-GSK-3β enhances the release of extracellular vesicles, such as exosomes, from cardiomyocytes under ischemic conditions. Exosomes contribute to cardiac regeneration through their content which can include mitogenic, angiogenic, anti-apoptotic and growth factors (Chistiakov et al., 2016). Many cardioprotective roles of CPC-derived exosomes have been reported which are mainly attributed to pro-angiogenic and cytoprotective miRNAs and proteins (Barile et al., 2017). Moreover, cardiomyocytes, fibroblasts and endothelial cells can communicate with each other by releasing the exosomes (Sluijter et al., 2014). Therefore, the release of growth factors from GSK3β-manipulated cardiomyocytes via exosomes in the ischemic heart is worth exploration in the field of cardiac repair.

We identified that the gain-of-GSK-3β function in human cardiomyocytes suppresses the level of secreted angpt2 post-hypoxia. Angpt2 plays an important role in vascular angiogenesis and enhances endothelial cell survival through stimulating PI3K-Akt signaling (Kim et al., 2000). It is also evident that human endometrium-derived stem cells transplantation facilitates post-MI cardiac repair due to an upregulation of pro-angiogenic molecules, including angpt2 (Fan et al., 2021). The role of angpt2 in cell proliferation or differentiation is unknown. However, it may have an effect on cardiomyocyte survival post-MI.

The dysregulated growth factors identified in GSK-3β overexpressing cardiomyocytes are known to regulate cardiac regeneration by modulating several cellular processes including cell proliferation, differentiation, cell survival and death, and CPC mobilization (Reboucas et al., 2016). Co-expression of VEGF and Angpt1 limits cardiomyocyte apoptosis and promotes cardiomyocyte proliferation and angiogenesis in the porcine ischemic heart (Taimeh et al., 2013; Tao et al., 2011). Similarly, myocardial injection of nanofibers linked with PDGF-BB in rats has shown protection against ischemia-induced cardiomyocyte apoptosis and preserved systolic function. PDGF-BB activates the Akt pathway through binding with cardiomyocyte PDGF receptor-β (PDGFR-β) *in vivo* (Hsieh et al., 2006). Consistently, a recent study has reported that PDGFR-β pathway promotes cardiomyocyte proliferation and heart regeneration (Yue et al., 2019). We found that both VEGF and PDGF were downregulated in GSK-3β overexpressing cardiomyocytes, particularly in the cells that had been exposed to hypoxia. Cardiomyocyte proliferation in the GSK-3β cKO heart may be induced by these growth factors through autocrine signaling. A study employing the miniswine model has shown that post-acute MI, cardiac injection of exogenous bFGF leads to an increased number of c-Kit- and 5-Bromo-2-deoxyuridine (BrdU)-positive cells, angiogenesis, myocardial regeneration, and, ultimately, improved cardiac function (Zhang et al., 2011). Our results are consistent with these findings where we identified a dysregulated bFGF in GSK-3β overexpressing cardiomyocytes post-hypoxia and an increased number of c-Kit-positive cells in the GSK-3β cKO hearts, post-MI. These findings suggest that GSK-3β deficient cardiomyocytes in the cKO heart likely secrete elevated levels of bFGF post-MI, which acts on resident CPCs through a paracrine mechanism.

Taken together, we have identified GSK-3β as a novel regulator of numerous growth factors in cardiomyocytes under stress conditions. These growth factors are critical for autocrine and paracrine signaling in the injured heart to stimulate cardiomyocyte and CPC proliferation, differentiation, and survival. Employing GSK-3β cKO mice and AC16 human cardiomyocytes, herein we demonstrate that GSK-3β inhibition could serve as a powerful tool to create a favorable environment for CPCs and cardiomyocytes to proliferate *in situ* and to accelerate the rate of cardiac repair in the ischemic heart. Furthermore, our findings suggest that targeting cardiomyocyte GSK-3β *ex vivo* could also be employed to synthesize important growth factors and, myocardial injection of these growth factors may potentially provide beneficial effects post-ischemia.

## Data Availability Statements

All data generated or analysed during this study are included in this article and additional information are available from the corresponding author on reasonable request.

## References

Ahmad F, Lal H, Zhou J, Vagnozzi RJ, Yu JE, Shang X, Woodgett JR, Gao E, Force T. 2014. Cardiomyocyte-specific deletion of gsk3alpha mitigates post-myocardial infarction remodeling, contractile dysfunction, and heart failure. J Am Coll Cardiol 64(7):696–706.

Ahmad F, Singh AP, Tomar D, Rahmani M, Zhang Q, Woodgett JR, Tilley DG, Lal H, Force T. 2019. Cardiomyocyte-GSK-3alpha promotes mPTP opening and heart failure in mice with chronic pressure overload. J Mol Cell Cardiol 130:65–75.

Ahmad F, Tomar D, Aryal ACS, Elmoselhi AB, Thomas M, Elrod JW, Tilley DG, Force T. 2020. Nicotinamide riboside kinase-2 alleviates ischemia-induced heart failure through P38 signaling. Biochim Biophys Acta Mol Basis Dis 1866(3):165609.

Ahmad F, Woodgett JR. 2020. Emerging roles of GSK-3alpha in pathophysiology: Emphasis on cardio-metabolic disorders. Biochim Biophys Acta Mol Cell Res 1867(2):118616.

Ahuja P, Sdek P, Maclellan WR. 2007. Cardiac myocyte cell cycle control in development, disease, and regeneration. Physiol Rev 87:521–544.

Barile L, Milano G, Vassalli G. 2017. Beneficial effects of exosomes secreted by cardiac-derived progenitor cells and other cell types in myocardial ischemia. Stem Cell Investig 4:93.

Bernstein ID, Andrews RG, Zsebo KM. 1991. Recombinant human stem cell factor enhances the formation of colonies by CD34+ and CD34+lin-cells, and the generation of colony-forming cell progeny from CD34+lin-cells cultured with interleukin-3, granulocyte colony-stimulating factor, or granulocyte-macrophage colony-stimulating factor. Blood 77(11):2316–2321.

Beurel E, Grieco SF, Jope RS. 2015. Glycogen synthase kinase-3 (GSK3): regulation, actions, and diseases. Pharmacol Ther 148:114–131.

Chistiakov DA, Orekhov AN, Bobryshev YV. 2016. Cardiac Extracellular Vesicles in Normal and Infarcted Heart. Int J Mol Sci 17(1).

Chute JP, Muramoto GG, Dressman HK, Wolfe G, Chao NJ, Lin S. 2006. Molecular profile and partial functional analysis of novel endothelial cell-derived growth factors that regulate hematopoiesis. Stem Cells 24(5):1315–1327.

Cohen P, Frame S. 2001. The renaissance of GSK-3. Nat Rev Mol Cell Biol 10:769–776.

Der Sarkissian S, Levesque T, Noiseux N. 2017. Optimizing stem cells for cardiac repair: Current status and new frontiers in regenerative cardiology. World J Stem Cells 9(1):9–25.

Doble BW, Patel S, Wood GA, Kockeritz LK, Woodgett JR. 2007. Functional redundancy of GSK-3alpha and GSK-3beta in Wnt/beta-catenin signaling shown by using an allelic series of embryonic stem cell lines. Dev Cell 12(6):957–971.

Fan X, He S, Song H, Yin W, Zhang J, Peng Z, Yang K, Zhai X, Zhao L, Gong H, Ping Y, Jiao X, Zhang S, Yan C, Wang H, Li RK, Xie J. 2021. Human endometrium-derived stem cell improves cardiac function after myocardial ischemic injury by enhancing angiogenesis and myocardial metabolism. Stem Cell Res Ther 12(1):344.

Force T, Woodgett JR. 2009. Unique and overlapping functions of GSK-3 isoforms in cell differentiation and proliferation and cardiovascular development. J Biol Chem 284(15):9643–9647.

Gao E, Lei YH, Shang X, Huang ZM, Zuo L, Boucher M, Fan Q, Chuprun JK, Ma XL, Koch WJ. 2010. A novel and efficient model of coronary artery ligation and myocardial infarction in the mouse. Circ Res 107:1445–1453.

Gupte M, Tumuluru S, Sui JY, Singh AP, Umbarkar P, Parikh SS, Ahmad F, Zhang Q, Force T, Lal H. 2018. Cardiomyocyte-specific deletion of GSK-3beta leads to cardiac dysfunction in a diet induced obesity model. Int J Cardiol.

He L, Nguyen NB, Ardehali R, Zhou B. 2020. Heart Regeneration by Endogenous Stem Cells and Cardiomyocyte Proliferation: Controversy, Fallacy, and Progress. Circulation 142(3):275–291.

Hodgkinson CP, Bareja A, Gomez JA, Dzau VJ. 2016. Emerging Concepts in Paracrine Mechanisms in Regenerative Cardiovascular Medicine and Biology. Circ Res 118(1):95–107.

Hsieh PC, Davis ME, Gannon J, MacGillivray C, Lee RT. 2006. Controlled delivery of PDGF-BB for myocardial protection using injectable self-assembling peptide nanofibers. J Clin Invest 116(1):237–248.

Iannolo G, Sciuto MR, Raffa GM, Pilato M, Conaldi PG. 2018. MiR34 inhibition induces human heart progenitor proliferation. Cell Death Dis 9(3):368.

Ji XK, Xie YK, Zhong JQ, Xu QG, Zeng QQ, Wang Y, Zhang QY, Shan YF. 2015. GSK-3beta suppresses the proliferation of rat hepatic oval cells through modulating Wnt/beta-catenin signaling pathway. Acta Pharmacol Sin 36(3):334–342.

Kaidanovich-Beilin O, Woodgett JR. 2011. GSK-3: Functional Insights from Cell Biology and Animal Models. Front Mol Neurosci 4:40.

Kerkela R, Kockeritz L, Macaulay K, Zhou J, Doble BW, Beahm C, Greytak S, Woulfe K, Trivedi CM, Woodgett JR, Epstein JA, Force T, Huggins GS. 2008. Deletion of GSK-3beta in mice leads to hypertrophic cardiomyopathy secondary to cardiomyoblast hyperproliferation. J Clin Invest 118(11):3609–3618.

Kim I, Kim JH, Moon SO, Kwak HJ, Kim NG, Koh GY. 2000. Angiopoietin-2 at high concentration can enhance endothelial cell survival through the phosphatidylinositol 3’-kinase/Akt signal transduction pathway. Oncogene 19(39):4549–4552.

Korur S, Huber RM, Sivasankaran B, Petrich M, Morin P, Jr., Hemmings BA, Merlo A, Lino MM. 2009. GSK3beta regulates differentiation and growth arrest in glioblastoma. PLoS One 4(10):e7443.

Laflamme MA, Murry CE. 2011. Heart regeneration. Nature 473(7347):326–335.

Lal H, Ahmad F, Woodgett J, Force T. 2015. The GSK-3 family as therapeutic target for myocardial diseases. Circ Res 116(1):138–149.

Lal H, Ahmad F, Zhou J, Yu JE, Vagnozzi RJ, Guo Y, Yu D, Tsai EJ, Woodgett J, Gao E, Force T. 2014. Cardiac fibroblast glycogen synthase kinase-3beta regulates ventricular remodeling and dysfunction in ischemic heart. Circulation 130(5):419–430.

Lal H, Zhou J, Ahmad F, Zaka R, Vagnozzi RJ, Decaul M, Woodgett J, Gao E, Force T. 2012. Glycogen synthase kinase-3alpha limits ischemic injury, cardiac rupture, post-myocardial infarction remodeling and death. Circulation 125(1):65–75.

Marian AJ, Braunwald E. 2017. Hypertrophic Cardiomyopathy: Genetics, Pathogenesis, Clinical Manifestations, Diagnosis, and Therapy. Circ Res 121(7):749–770.

Patel S, Doble BW, MacAulay K, Sinclair EM, Drucker DJ, Woodgett JR. 2008. Tissue-specific role of glycogen synthase kinase 3beta in glucose homeostasis and insulin action. Mol Cell Biol 28(20):6314–6328.

Porrello ER, Mahmoud AI, Simpson E, Hill JA, Richardson JA, Olson EN, Sadek HA. 2011. Transient regenerative potential of the neonatal mouse heart. Science 331(6020):1078–1080.

Reboucas JS, Santos-Magalhaes NS, Formiga FR. 2016. Cardiac Regeneration using Growth Factors: Advances and Challenges. Arq Bras Cardiol 107(3):271–275.

Savarese G, Lund LH. 2017. Global Public Health Burden of Heart Failure. Card Fail Rev 3(1):7–11.

Segers VF, Lee RT. 2008. Stem-cell therapy for cardiac disease. Nature 451(7181):937–942.

Singh AP, Umbarkar P, Guo Y, Force T, Gupte M, Lal H. 2019. Inhibition of GSK-3 to induce cardiomyocyte proliferation: a recipe for in situ cardiac regeneration. Cardiovasc Res 115(1):20–30.

Sluijter JP, Verhage V, Deddens JC, van den Akker F, Doevendans PA. 2014. Microvesicles and exosomes for intracardiac communication. Cardiovasc Res 102(2):302–311.

Taimeh Z, Loughran J, Birks EJ, Bolli R. 2013. Vascular endothelial growth factor in heart failure. Nat Rev Cardiol 10(9):519–530.

Takahashi-Yanaga F. 2018. Roles of Glycogen Synthase Kinase-3 (GSK-3) in Cardiac Development and Heart Disease. J UOEH 40(2):147–156.

Tao Z, Chen B, Tan X, Zhao Y, Wang L, Zhu T, Cao K, Yang Z, Kan YW, Su H. 2011. Coexpression of VEGF and angiopoietin-1 promotes angiogenesis and cardiomyocyte proliferation reduces apoptosis in porcine myocardial infarction (MI) heart. Proc Natl Acad Sci U S A 108(5):2064–2069.

Urbanek K, Rota M, Cascapera S, Bearzi C, Nascimbene A, De Angelis A, Hosoda T, Chimenti S, Baker M, Limana F, Nurzynska D, Torella D, Rotatori F, Rastaldo R, Musso E, Quaini F, Leri A, Kajstura J, Anversa P. 2005a. Cardiac stem cells possess growth factor-receptor systems that after activation regenerate the infarcted myocardium, improving ventricular function and long-term survival. Circ Res 97(7):663–673.

Urbanek K, Torella D, Sheikh F, De Angelis A, Nurzynska D, Silvestri F, Beltrami CA, Bussani R, Beltrami AP, Quaini F, Bolli R, Leri A, Kajstura J, Anversa P. 2005b. Myocardial regeneration by activation of multipotent cardiac stem cells in ischemic heart failure. Proc Natl Acad Sci U S A 102:8692–8697.

Williams DE, Foxworthe D, Teepe M, Lyman SD, Anderson D, Eisenman J. 1992. Recombinant murine steel factor stimulates in vitro production of granulocyte-macrophage progenitor cells. J Cell Biochem 50(3):221–226.

Witman N, Zhou C, Grote Beverborg N, Sahara M, Chien KR. 2020. Cardiac progenitors and paracrine mediators in cardiogenesis and heart regeneration. Semin Cell Dev Biol 100:29–51.

Woldbaek PR, Hoen IB, Christensen G, Tonnessen T. 2002. Gene expression of colony-stimulating factors and stem cell factor after myocardial infarction in the mouse. Acta Physiol Scand 175(3):173–181.

Woulfe KC, Gao E, Lal H, Harris D, Fan Q, Vagnozzi R, DeCaul M, Shang X, Patel S, Woodgett JR, Force T, Zhou J. 2010. Glycogen synthase kinase-3beta regulates post-myocardial infarction remodeling and stress-induced cardiomyocyte proliferation in vivo. Circ Res 106(10):1635–1645.

Yoon YWA, Heyd L, Park JS, Tkebuchava T, Kusano K, Hanley A, Scadova H, Qin G, Cha DH, Johnson KL, Aikawa R, Asahara T, Losordo DW. 2005. Clonally expanded novel multipotent stem cells from human bone marrow regenerate myocardium after myocardial infarction. J Clin Inv 115:326–338.

Yu H, Wang Z. 2019. Cardiomyocyte-Derived Exosomes: Biological Functions and Potential Therapeutic Implications. Front Physiol 10:1049.

Yue Z, Chen J, Lian H, Pei J, Li Y, Chen X, Song S, Xia J, Zhou B, Feng J, Zhang X, Hu S, Nie Y. 2019. PDGFR-beta Signaling Regulates Cardiomyocyte Proliferation and Myocardial Regeneration. Cell Rep 28(4):966–978 e964.

Zhang LX, DeNicola M, Qin X, Du J, Ma J, Tina Zhao Y, Zhuang S, Liu PY, Wei L, Qin G, Tang Y, Zhao TC. 2014. Specific inhibition of HDAC4 in cardiac progenitor cells enhances myocardial repairs. Am J Physiol Cell Physiol 307(4):C358–372.

Zhang YH, Zhang GW, Gu TX, Li-Ling J, Wen T, Zhao Y, Wang C, Fang Q, Yu L, Liu B. 2011. Exogenous basic fibroblast growth factor promotes cardiac stem cell-mediated myocardial regeneration after miniswine acute myocardial infarction. Coron Artery Dis 22(4):279–285.

Zhou J, Ahmad F, Lal H, Force T. 2016a. Response by Zhou et al to Letter Regarding Article, “Loss of Adult Cardiac Myocyte GSK-3 Leads to Mitotic Catastrophe Resulting in Fatal Dilated Cardiomyopathy”. Circ Res 119(2):e29–e30.

Zhou J, Ahmad F, Parikh S, Hoffman NE, Rajan S, Verma VK, Song J, Yuan A, Shanmughapriya S, Guo Y, Gao E, Koch W, Woodgett JR, Madesh M, Kishore R, Lal H, Force T. 2016b. Loss of Adult Cardiac Myocyte GSK-3 Leads to Mitotic Catastrophe Resulting in Fatal Dilated Cardiomyopathy. Circ Res 118(8):1208–1222.

