## Supplemental Figure for "Cardiomyocyte GSK-3β deficiency induces cardiac progenitor cell proliferation in the ischemic heart through paracrine mechanisms"

### Suppl. Figure 1

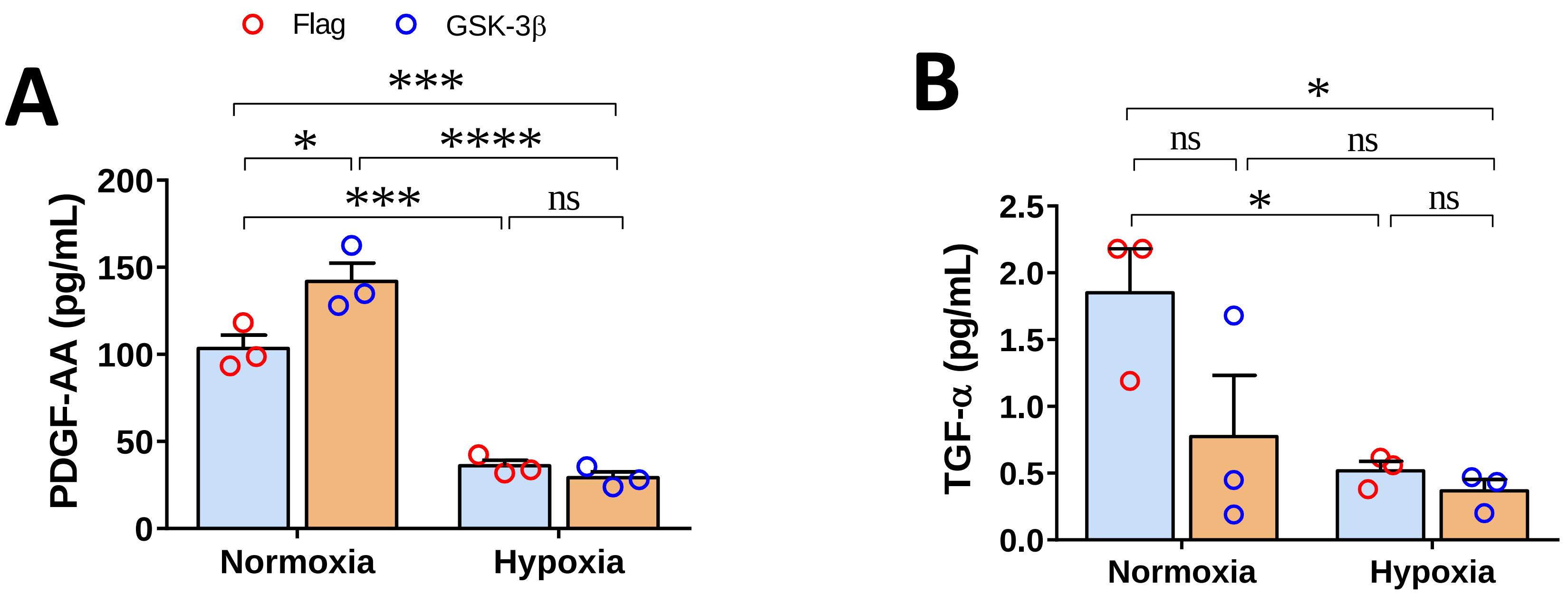


**Supplemental figure 1; Role of cardiomyocyte-GSK-3β in growth factor expression and secretion post-hypoxia:** Bar diagrams show significantly lower levels of (**A**) platelet-derive growth factor-AA (PDGF-AA), and (**B**) transforming growth factor-α (TGF-α) in the culture supernatant of control and GSK-3β overexpressing cardiomyocyte post-hypoxia compared to normoxia group. ns= non-significant; * $P$<0.05 , *** $P$<0.001, **** $P$<0.0001.

### Suppl. Figure 2

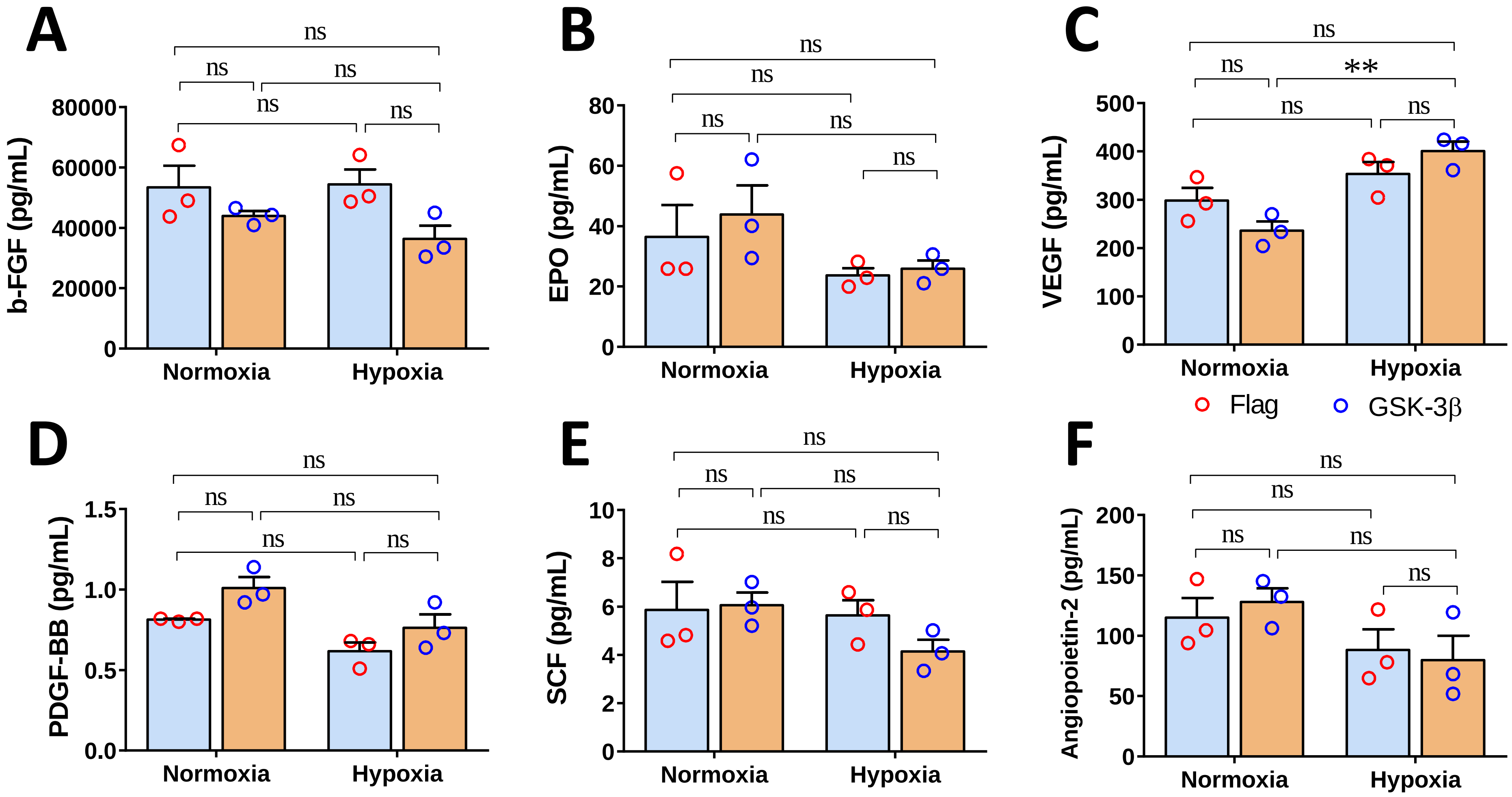


**Supplemental figure 2; Growth factor expression level in GSK-3β overexpressing cardiomyocyte post-hypoxia:** Bar diagrams show a comparable levels of (**A**) basic-fibroblast growth factor (b-FGF) (**B**) erythropoietin (EPO), (**C**) vascular endothelial growth factor (VEGF), (**D**) platelet-derive growth factor-BB (PDGF-BB), (**E**) stem cell factor (SCF), and (**F**) angiopoietin-2 in the GSK-3β overexpressing compared to control cardiomyocytes under both normoxia and hypoxia conditions. ns= non-significant.

### Suppl. Figure 3

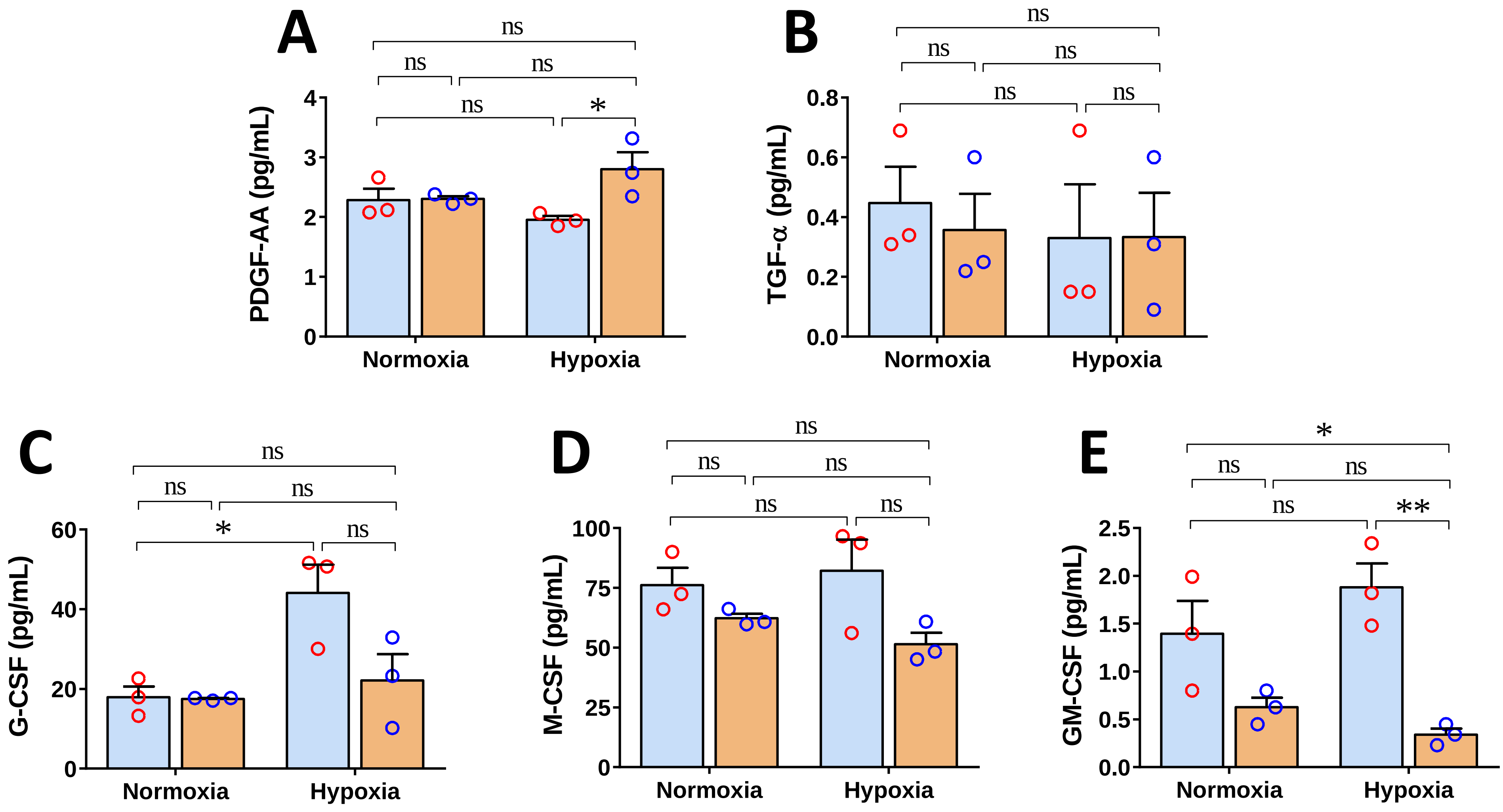


**Supplemental figure 3; Role of cardiomyocyte-GSK-3β in growth factor expression post-hypoxia:** Scatter dot plots show (**A**) a significantly higher level of platelet-derive growth factor-AA (PDGF-AA), (**B**) comparable transforming growth factor-α (TGF-α), and a trend of lower levels of (**C**) granulocyte-colony stimulating factor (G-CSF), (**D**) macrophage-CSF (M-CSF), and (**E**) granulocyte macrophage- CSF (GM-CSF) in GSK-3β overexpressing cardiomyocytes post-hypoxia. ns= non-significant; * *P*<0.05 , ** *P*<0.005.
